# Inhibitory activities of monoclonal antibodies against *Staphylococcus aureus* Clumping factor A

**DOI:** 10.1101/2025.05.21.655433

**Authors:** Biswarup Banerjee, Carla Emolo, Miaomiao Shi, Abrar Al Fardan, Tonu Pius, Molly McAdow, Olaf Schneewind, Dominique Missiakas

## Abstract

*Staphylococcus aureus* infection is a frequent cause of sepsis in humans, a disease associated with high mortality and without specific intervention. Clumping factor A (ClfA) displayed on the bacterial surface plays a key role in promoting *S. aureus* replication during invasive disease. Decades of research have pointed to a wide array of ligands engaged by ClfA. The sum of these interactions supports the unique ability of this pathogen to survive and replicate in the blood stream. One such ligand is fibrin. ClfA acts as the key agglutinating factor of *S. aureus* by promoting the shielding of bacteria in fibrin cables and their physical escape from phagocytes. Here, we compare a series of monoclonal antibodies elicited against the ligand binding domain of ClfA following immunization of mice. We analyze these antibodies for their ability to neutralize ClfA interactions and to promote the uptake of staphylococci in whole blood. We find that while all the antibodies promoted opsonophagocytic uptake, only those that also inhibited ClfA interactions with ligands, reduced bacterial burdens in animals following blood stream challenge with *S. aureus*.

**IMPORTANCE:** Antibody-based approaches to fight bacterial pathogens have been modeled on toxin-producing or encapsulated pathogens for which correlates of protection can be reduced to measuring antibody neutralization or complement-fixing activities using tissue cultured cells. Such approaches have failed against *Staphylococcus aureus* raising uncertainty about the value of antibodies. Here, we use a series of mouse monoclonal antibodies directed against ClfA, a surface protein that allows *S. aureus* to thrive in the blood stream, to query how antibodies may be exploited against a pathogen endowed with a formidable array of virulence factors.

## INTRODUCTION

*Staphylococcus aureus* is a human pathogen that resides on the skin and nares but has the unique ability to replicate in the blood stream to cause mild to life threatening infections (1, 2). Spillover to animals in close contact to humans and genetic adaptation also result in anthropogenic infections; as an example, *S. aureus* is a major cause of bovine mastitis in dairy cattle (3–5). *S. aureus* relies on surface proteins to breach host barriers and escape host defenses (6, 7). In *S. aureus*, most surface proteins are covalently linked to peptidoglycan by sortase enzymes (8, 9). Mutants lacking the housekeeping sortase A enzyme have been found to be avirulent in animal models (10, 11). Although sortase A anchors about 24 proteins on the bacterial surface, the exact count varying by strain (8, 9), Clumping factor A (ClfA) has been shown to be critical for infection (12). Experiments have shown that animals are more likely to survive an intravenous *S. aureus* infection when challenged with *clfA* mutant strains (11, 13). ClfA mediates substrate interactions that enhance *S. aureus* adhesion to vascular epithelia and catheter-induced thrombi, facilitate *S. aureus* escape from neutrophils, and promote invasion into tissues including joints (14–18). Thus, ClfA is important in promoting *S. aureus* bacteremia, septic death, septic arthritis, endocarditis, and abscess formation in deep-seated organs (11, 13, 19–22). As a result, ClfA has emerged as a prime antigen target for the development of immune-based therapies in both humans and domestic animals, the latter in an attempt to reduce the incidence of bovine mastitis (19, 23–32).

ClfA is a member of the Clf-Sdr-FnBP sub-family of microbial surface components recognizing adhesive matrix molecules (MSCRAMMs) (12). MSCRAMMs are characterized by the presence of two adjacent IgG-like subdomains responsible for substrate binding (12). *Staphylococcus epidermidis* SdrG interaction with the β-chain of fibrinogen exemplifies a binding mechanism described as ‘dock-lock-latch’ (DLL) and involves a trench between the two IgG-like subdomains (33, 34). In ClfA, the IgG-like subdomains are encompassed by the N2-N3 segment in the N-terminal A domain (ClfA-A) (35) (Fig. 1A); the hydrophobic trench between N2 and N3 captures the last 17-residues of the γ-chain of fibrinogen and a conformational change at the C-terminus of N3 locks the peptide in place (36, 37). Fibrinogen is a glycoprotein (∼ 340 kDa) composed of three pairs of covalently linked Aα-, Bβ-, and γ-chains with the letters A and B designating the fibrinopeptides released by thrombin cleavage (38). In the soluble dimer, the N-termini of all six chains form the central E domain and each trimer extends at opposite ends through coiled-coil segments to form two symmetrical D domains composed mainly by the C-termini of the Bβ- and γ-chains (38). This symmetry accounts for the clumping activity of fibrinogen when added to *S. aureus* bacterial suspensions and the naming of ClfA (36, 39). But two decades of research have unveiled additional ClfA ligands. Recombinant ClfA has been shown to bind Complement factor I, bovine Annexin 2, and secreted *S. aureus* von Willebrand binding protein (vWbp), highlighting the pleiotropic roles of ClfA during infection (15, 17, 18, 40). Importantly, ClfA also promotes agglutination, the interaction of bacteria with fibrin, the insoluble form of fibrinogen produced during blood clotting, a process hijacked by the secreted coagulases of *S. aureus*, coagulase (Coa) and vWbp (13, 41).

**Fig. 1.**
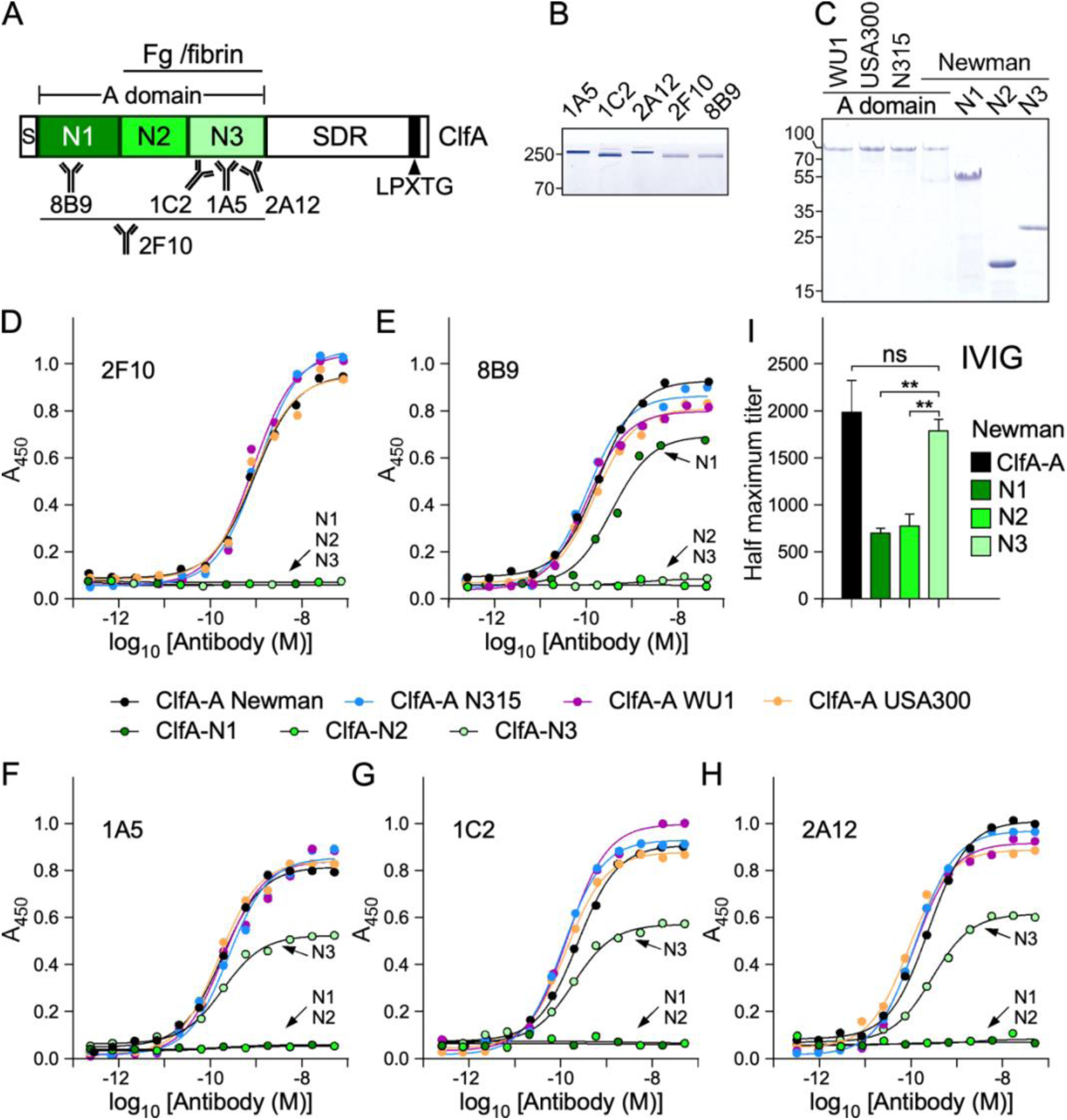
Characterization of anti-ClfA mAbs. (A) Domain organization of ClfA depicting the signal sequence (S), N1, N2, N3 subdomains, A domain (ClfA-A), serine aspartic repeat region (SDR), LPXTG motif for attachment to peptidoglycan by sortase A (black rectangle). N2-N3 segment involved in fibrinogen (Fg)/fibrin binding and binding sites of mAbs deduced from this study are shown. (B, C) Coomassie stained gels of purified mAbs (non-reducing conditions) (B) and ClfA domains and subdomains from various strains (C). Numbers to the left of gels indicate molecular weight markers in kDa. (D-H) Representative ELISA showing mAb binding to ClfA-A Newman, N315, WU1, USA300 as well as isolated, ClfA-N1, -N2 or -N3 subdomains from strain Newman. Antibodies are presented as follows: 2F10 (D), 8B9 (E), 1A5 (F), 1C2 (G), 2A12 (H). Bound antibodies were detected with polyclonal anti-mouse HRP-conjugated secondary antibody and reported as absorbance at 450 nm (A_450_). Association constants and comparative analyses are shown in Table 1 and Supplementary Table 1. (I) Relative abundance of Newman ClfA-A, N1, N2 and N3 antibodies in IVIG. Data was analyzed with GraphPad Prism 10 using non-linear fit (least squared) regression. (**, *P* < 0.01; ns: not significant). All experiments were performed at least twice.

**Table 1.** Interactions between mAbs and various ClfA antigens as determined by ELISA.

| Antibody/Antigen | $EC_{50} (\times 10^{-9} \text{M}) \pm \text{SEM} (\times 10^{-9})$ | | | | | | |
| --- | --- | --- | --- | --- | --- | --- | --- |
|  | N1 | N2 | N3 | ClfA-A<br>Newman | ClfA-A<br>N315 | ClfA-A<br>WU1 | ClfA-A<br>USA300 |
| **1C2-mIgG1** | / | / | 1.8 ± 1.8 | 2.2 ± 1.1 | 1.1 ± 1.6 | 1.3 ± 1.5 | 1.8 ± 1.3 |
| **2F10-mIgG1** | / | / | / | 1.3 ± 0.3 | 1.2 ± 0.03 | 1.5 ± 0.02 | 1.5 ± 0.01 |
| **8B9-mIgG1** | 2.6 ± 1.8 | / | / | 2 ± 1.5 | 1.2 ± 1.5 | 1.1 ± 1.5 | 1.8 ± 1.6 |
| **mIgG1** | / | / | / | / | / | / | / |
| **1A5-mIgG2a** | / | / | 1.9 ± 1.5 | 1.9 ± 1.6 | 2.6 ± .73 | 1.6 ± 1.6 | 1.9 ± 1.6 |
| **mIgG2a** | / | / | / | / | / | / | / |
| **2A12-mIgG2b** | / | / | 2.7 ± 0.6 | 2.2 ± 1.2 | 1.7 ± 1.6 | 1.1 ± 1.7 | 1 ± 1.8 |
| **mIgG2b** | / | / | / | / | / | / | / |

Here, we isolated a series of monoclonal antibodies (mAbs) by immunizing mice with ClfA-A and characterized these antibodies for their ability to block *S. aureus* interactions with host ligands, promote opsonophagocytosis of bacteria in freshly drawn blood, and protect mice in a model of blood stream infection. Our findings suggest that the ability to block ClfA interactions with ligands, but not the simple opsonophagocytic activity of antibodies, is critical for protecting animals following a blood stream challenge. Thus, carefully designed antibodies targeting ClfA have the potential to reduce the severity of invasive disease associated with *S. aureus* bacteremia.

## RESULTS

### Characterization of mAbs raised against ClfA-A

mAbs were raised by immunizing mice with the A domain of *S. aureus* Newman ClfA, ClfA-A (42). The amino acid sequence of this antigen, residues 40-559, is shown in Fig. S1. Of note, the only known structure of ClfA corresponds to the N2-N3 segment (residues 221-559) (35). The antigen used here was produced in *Escherichia coli* without its cleavable signal sequence (residues 1-39) or repeats of serine aspartic (SD) amino acids (residues 560-668) that otherwise tether the A domain to the C-terminal sorting signal (residues 669-933) for attachment to the cell wall (Fig. 1A). In *S. aureus*, the SD repeats are extensively modified with N-acetyl-glucosamine and are not thought to bind any ligand (43, 44). Five antibodies reactive to ClfA-A were identified and purified (Fig. 1B) and characterized for their specificity and affinity to the full-length A domain as well as the separated N1, N2 and N3 subdomains of strain Newman (Fig. 1C). The various antigens were coated on 96-well plates and subjected to enzyme-linked immunosorbent assay (ELISA) using serial dilutions of each mAb (Fig. 1D-H). mAb 2F10 was found to interact only with the full-length A domain but not with the isolated N1, N2 or N3 subdomains (Fig. 1D). The remaining antibodies bound the isolated N1 subdomain, mAb 8B9 (Fig. 1E), and the isolated N3 subdomain, mAbs 1A5, 1C2, 2A12, respectively (Fig. 1FGH; Table 1). When binding was detected, the half-maximal effective concentration (EC_50_) values were not found to be statistically significant in any pairwise comparison (Supplementary Table 1) meaning that all mAbs bound the ligands with similar affinity.

The amino acid sequence of ClfA varies between sequenced isolates, a challenge noted earlier in the design a human vaccine including ClfA (45, 46) and the development of mAbs (28, 47). To assess the specificity of mAbs isolated in our studies, the A domain was also produced and purified using sequences of strains USA300 (USA MRSA; (48)), N315 (Japanese MRSA; (49)), and WU1 (Fig. 1C; Fig. S1). The ClfA-A sequences of strain Newman and USA300 are identical with the exception of one substitution in the N1 subdomain. Greater variability exists between the ClfA-A sequences of strains N315 and WU1. WU1 is a mouse-adapted strain used in our laboratory to study *S. aureus* colonization (50, 51). ELISA experiments revealed that each of the five mAbs interacted similarly with the four ClfA-A variants regardless of their allelic differences (Fig. 1D-H; Table 1, Supplementary Table 1). ELISA was also used to determine the isotype of the five mAbs and revealed that 1C2, 2F10 and 8B9 belong to the mIgG1 subclass, 1A5 belongs to the mIgG2a subclass, and 2A12 to the mIgG2b subclass (Fig. S2). Since three mAbs interacted with N3, a competition ELISA was performed with HRP labeled 2A12 and unlabeled 1A5, 1C2 and 2A12. This approach revealed that 1A5 and 2A12 may bind the same epitope (Fig. S3).

Three of our mAbs recognized the N3 subdomain, a property also shared with Tefibazumab, a humanized antibody derived from the mouse hybridoma MAb 12-9 (47). Our hybridomas were generated using the same antigen, mouse line and immunization protocol (47). We wondered if this approach may have introduced some bias toward N3 recognition. To examine this possibility, commercially available intravenous immunoglobulin (IVIG) was assessed for the presence of antibodies against ClfA-A, as well as the isolated N1, N2 and N3 subdomains. This analysis revealed that the N3 immunodominance is not restricted to mice as ClfA antibodies in human preferentially bind N3 (Fig. 1I).

### Assessing mAbs for the disruption of ClfA interaction with human fibrinogen and fibrin

ClfA is a conserved protein that mediates *S. aureus* interaction with fibrinogen. Despite their sequence differences, we observed that all ClfA-A variants used in this study interacted with similar affinity with human fibrinogen (Fig. 2A; Table 2). Next, fibrinogen was mixed with serial dilutions of mAbs before addition to immobilized ClfA-A. This approach showed that all the mAbs, with the exception of 8B9 (with N1 specificity) inhibited ClfA-A interaction with fibrinogen in a concentration dependent manner and regardless of differences in amino acid sequences (Table 3); this competition was also depicted as the fraction of fibrinogen that remains bound to ClfA-A Newman in the presence of 25 nM of mAbs (Fig. 2B).

**Fig. 2.**
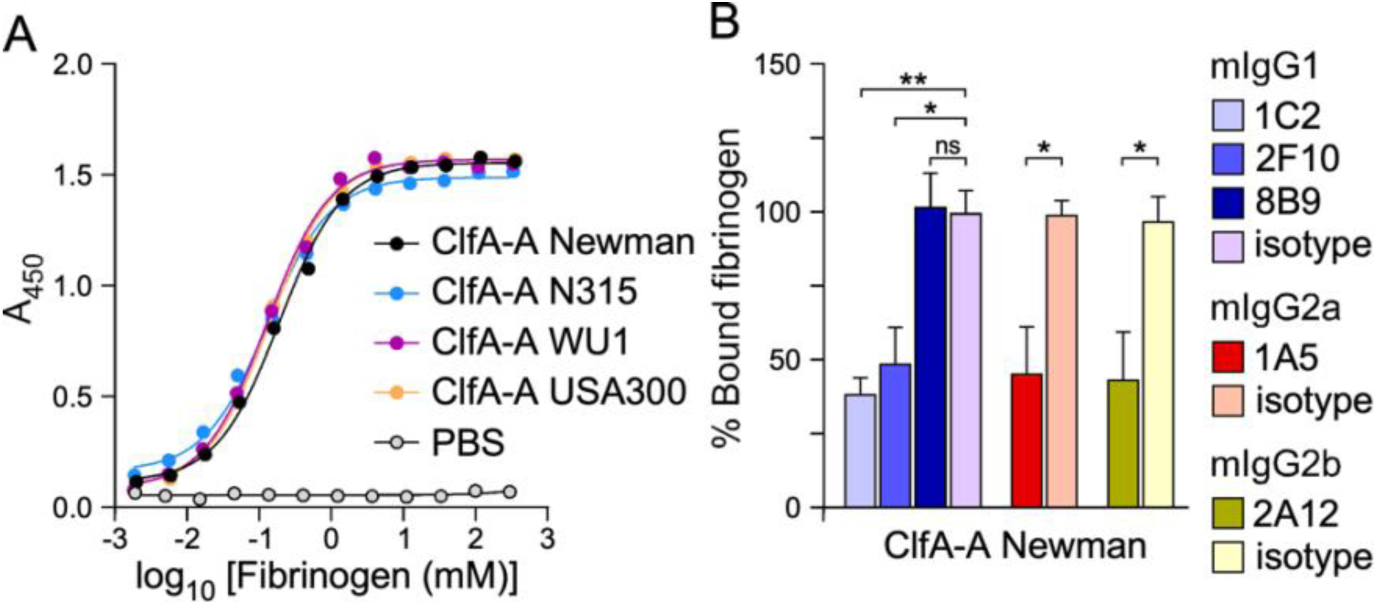
mAb inhibition of ClfA-A interaction with soluble fibrinogen. (A) Association of soluble fibrinogen with immobilized ClfA-A proteins from various strains. Bound fibrinogen was detected with polyclonal anti-human fibrinogen-HRP and reported as A_450_. Half-maximal effective concentrations (EC_50_) are reported in Table 2. (B) The ability of ClfA antibodies to compete for ClfA-A/fibrinogen interactions was measured using ELISA. Half-maximal inhibitory concentrations (IC_50_) are reported in Table 3. This panel shows the % of fibrinogen remaining bound to ClfA-A in the presence of 25 nM of antibodies. A_450_ values of bound fibrinogen in the presence of isotype control antibodies were set at 100%. Data are represented as mean ± SEM and statistical significance was calculated using One-way ANOVA with Tukey’s multiple comparison (**, *P* < 0.01; *, *P* < 0.05; ns: not significant). All experiments were performed at least twice.

**Table 2.** Interactions between various host and bacterial ligands.

| Human/Bacterial ligand | *EC*<sub>50</sub> (×10<sup>-6</sup> M) ± SEM (×10<sup>-6</sup>)* |  |  |  |  |  |
| --- | --- | --- | --- | --- | --- | --- |
|  | ClfA-A Newman | ClfA-A N315 | ClfA-A WU1 | ClfA-A USA300 | vWbp Newman | vWbp + ClfA-A Newman |
| **Fibrinogen** | 1.3±0.01 | 1.3±0.08 | 1.1±0.02 | 1±0.09 | ND | ND |
| **vWF** | / | ND | ND | ND | 14.5±1.69<sup>#</sup> | 3.5±0.35<sup>#</sup> |
| **Complement factor I** | 1.8±0.3 | ND | ND | ND | ND | ND |

**Table 3.** Inhibition of ClfA-A interaction with various ligands by mAbs.

| Human ligand | $IC_{50} (\times 10^{-10} M) \pm SEM (\times 10^{-10})$ | | | | | |
| --- | --- | --- | --- | --- | --- | --- |
|  | Fibrinogen |  |  |  | vWF | Complement factor I |
| Bacterial ligand | ClfA-A Newman | ClfA-A N315 | ClfA-A WU1 | ClfA-A USA300 | ClfA-A + vWbp Newman | ClfA-A Newman |
| 1C2-mIgG1 | 1.8±0.4 | 1.3±0.7 | 1.6±0.4 | 2±1.3 | 2.5±0.3 | / |
| 2F10-mIgG1 | 1.2±0.7 | 1.1±0.3 | 1±0.6 | 1.1±0.4 | 5.7±0.2 | / |
| 8B9-mIgG1 | / | / | / | / | / | / |
| mIgG1 | / | / | / | / | / | / |
| 1A5-mIgG2a | 1±0.58 | 1±1 | 1±0.7 | 1.2±1.1 | 8.1±0.43 | / |
| mIgG2a | / | / | / | / | / | / |
| 2A12-mIgG2b | 1.3±0.3 | 1.9±0.2 | 1.3±1 | 1.1±0.7 | 3.6±0.2 | / |
| mIgG2b | / | / | / | / | / | / |

Although, when mixed with purified fibrinogen, *S. aureus* cells form small clumps, when inoculated in anti-coagulated blood or plasma, *S. aureus* bacteria agglutinate (52). Both clumping and agglutination are ClfA-dependent (13, 36, 39). *S. aureus* agglutination can be visualized as the assembly of large aggregates in a network of fibrin, a reaction that requires the host factors prothrombin and fibrinogen (52), as well as ClfA and secreted Coa and vWbp, the factors that bind and activate prothrombin (13, 53). Thus, agglutination represents a more physiological host-pathogen interaction encountered during bacteremia. We asked if our candidate mAbs could reduce *S. aureus* Newman agglutination in mouse plasma. Bacteria were fluorescently stained with SYTO-9 and incubated with EDTA-plasma with mAbs or isotype control antibodies on a glass microscope slide for 30 minutes. Representative images shown in Fig. 3A were used to quantify the size of agglutinates under the various treatments (Fig. 3B). This approach revealed that all N3 (1A5, 1C2 and 2A12) and A domain-only binding (2F10) mAbs significantly reduced bacterial agglutination in plasma compared to cognate isotypes, but no reduction was observed for N1 binding, mAb 8B9 (Fig. 3). Newman and an isogenic *clfA* mutant with no additional treatments were used as positive and negative controls, respectively (Fig. 3). Together, these experiments demonstrate that mAbs that prevent ClfA-fibrinogen interaction also block *S. aureus* agglutination in plasma.

**Fig. 3.**
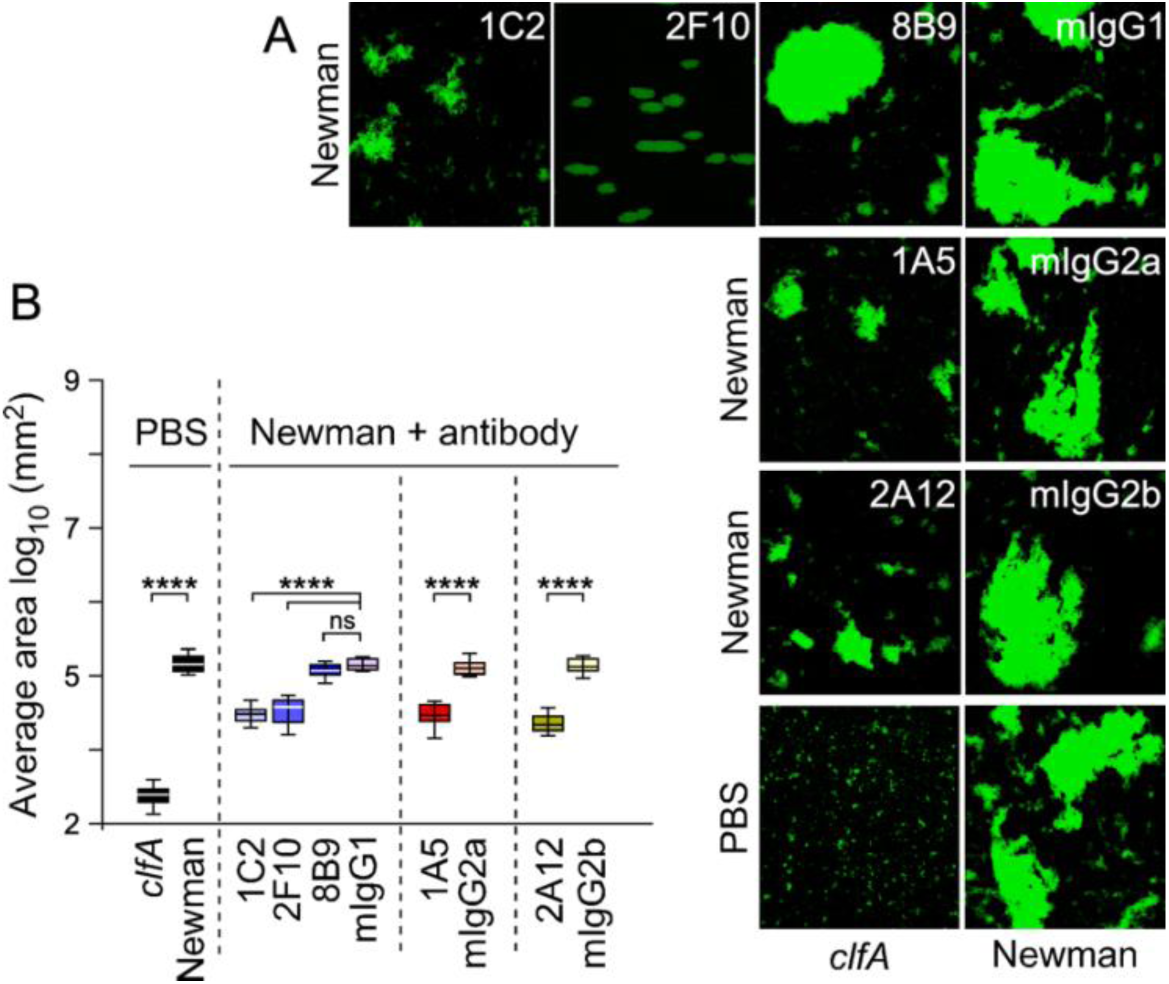
Staphylococcal agglutination in anti-coagulated mouse plasma. (A) Representative images of SYTO-9-stained Newman bacteria agglutinated in mouse plasma in the presence of isotype control or test antibodies that were obtained using an inverted fluorescent microscope with a 20× objective. (B) Box and whisker plot representation of agglutination areas with anti-coagulated mouse plasma collected from 10 fields of microscopic view. Statistical significance was assessed in pairwise comparison (wild type Newman versus *clfA* mutant; isotype control versus test antibody) using Brown-Forsythe and Welch ANOVA followed by Dunnett’s T3 correction for multiple comparisons (****, *P* < 0.0001; ns: not significant). All experiments were performed at least twice.

### mAbs reduce *S. aureus* adhesion by impeding ClfA-vWbp-vWF tripartite interaction

In the vasculature, ClfA promotes *S. aureus* adhesion via a fibrinogen bridge between the bacterial cell and the α_V_β_3_ integrin on the endothelial cell surface (54). Thus, antibodies blocking fibrinogen interactions would also prevent such adhesion. Genetic and biochemical studies have also demonstrated that secreted vWbp interacts with ClfA to promote *S. aureus* adhesion under shear blood flow in a fibrinogen-independent manner (17, 18, 55). Shear stress unfolds secreted von Willebrand Factor (vWF) unmasking sites for binding to the α_V_β_3_ integrin on endothelial cells of blood vessel walls (56). This unfolding also unmasks sites for both platelet and *S. aureus* adhesion (17). Like platelets, *S. aureus* binds directly to the A1 domain of vWF, utilizing an ultra-strong bond between ClfA and vWbp as measured using atomic force microscopy (17, 57). The weak interaction between purified vWbp and vWF was recapitulated in our ELISA and found to be significantly increased upon addition of ClfA-A (Table 2) confirming the earlier report (58). Addition of mAbs, with the exception of N1 binding mAb 8B9, disrupted the ClfA-A-mediated interaction between vWbp and vWF, in a dose-dependent manner (Table 3). To more directly assess the impact of mAbs on *S. aureus* adhesion under flow, fluorescently labeled bacteria were perfused at 1ml/min over human umbilical vein endothelial cells (HUVECs) seeded in a chambered coverslip. HUVECs were activated with a Ca^2+^-ionophore to cause the release of vWF and bacterial attachment captured with a fluorescence microscope in the presence of anti-ClfA mAbs or isotype control. Representative images are shown in Fig. 4A. Quantification of bound bacteria revealed that all mAbs with the exception of 8B9 reduced *S. aureus* adhesion (Fig. 4B). Newman and isogenic *clfA* and *vwb* mutants with no additional treatments served as positive and negative controls (Fig. 4). Together, the results indicate that antibodies that inhibit fibrin and fibrinogen interactions, also have the potential to block ClfA-mediated interactions critical for bacterial adhesion in the vasculature.

**Fig. 4.**
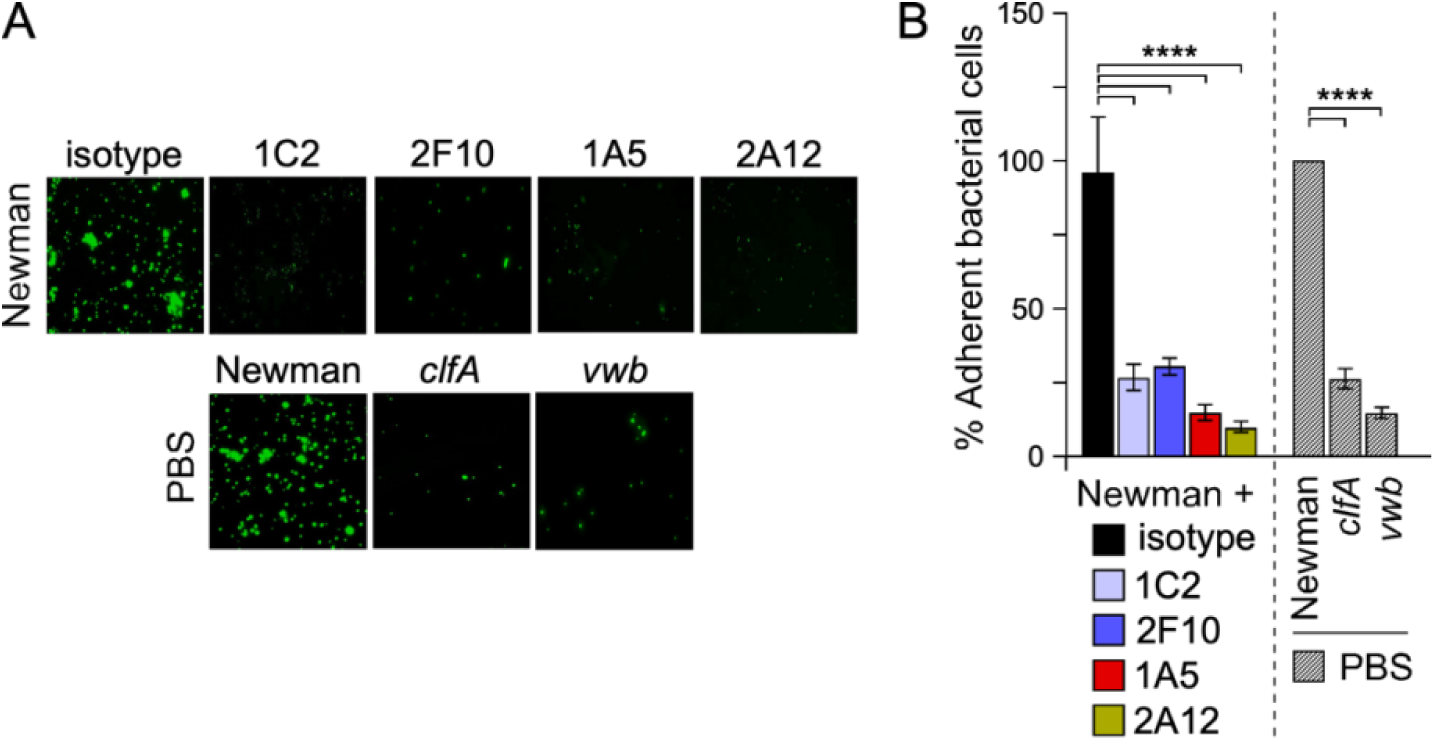
Staphylococcal adhesion to HUVECs under flow. (A) Representative images of SYTO-9-stained Newman bacteria that adhered to HUVEC cells under flow in iBidi microslide. Images were captured using an inverted fluorescent microscope with a 40× objective. (B) Quantification of adhesion as shown in A was performed by scanning fluorescence signals from 10 fields of microscopic view. Adhesion is plotted as percentage with 100 percent representing signals obtained for wild type bacteria with no additional treatment (Newman/PBS). Data are displayed as mean ± SEM and statistical significance was calculated using One-way ANOVA with Tukey’s multiple comparison (****, *P* < 0.0001). All experiments were performed twice for reproducibility.

### mAbs promote opsonophagocytic killing of *S. aureus* in mouse blood

In addition to inhibiting ClfA-ligand interactions, we would expect that antibodies targeting a surface antigen would promote bacterial uptake and killing by phagocytic cells. We used freshly drawn anticoagulated mouse blood supplemented with candidate or isotype control antibodies to evaluate the opsonophagocytic killing potential of mAbs. Since the agglutinating activity of ClfA shields staphylococci in fibrin cables, this approach is deemed more relevant than using purified neutrophils or differentiated HL-60 cells in an uptake assay (59). Bacteria were incubated in freshly drawn blood with antibodies for 30 min. Longer incubations lead to the deterioration of samples with extensive lysis of host cells. At the end of this incubation and before plating for colony forming enumeration (CFU), samples were treated with streptokinase and nucleases to liberate bacteria trapped in fibrin clots or neutrophil extracellular traps, and with saponin to release intracellular staphylococci not killed by immune cells. Counts at 30 min were reported as % with 100% representing CFU counts at time 0. In one set of experiments, the assay was performed using blood pre-treated for 10 min with cytochalasin D (CD), an inhibitor of actin polymerization that prevents phagocytosis by host cells (Fig. 5A). This was done to document if antibodies promoted phagocytic uptake of bacteria. To query the requirement for complement, in a second set of experiments, blood was preincubated with Cobra Venom Factor (CVF), a protein analogue of complement component C3 that continuously activates complement resulting in its depletion (60) (Fig. 5B). We observed an approximately 50% reduction in bacterial counts in samples treated with the anti-ClfA mAbs compared to the respective isotype controls (Fig. 5AB; left panels). No statistical differences were observed between candidate mAbs. Addition of CD eliminated any protective activity mediated by mAbs (Fig. 5A). Addition of CVF had no impact on the activity of 1C2/2F10/8B9-mIgG1 and 2A12-mIgG2b, but preempted any protection mediated by 1A5-mIgG2a (Fig. 5B). These results agree with the notion that mIgG1 and mIgG2a display negligible and high interactions with C1q, respectively (Fig. 5C) (61, 62), a behavior that is corroborated when assessing mAb-mediated C1q recruitment on the surface of *S. aureus* (Fig. 5D). Thus, the loss of killing activity by 1A5-mIgG2a upon depletion of complement conforms with the intrinsic ability of this antibody to interact with C1q. But the finding that 2A12-mIgG2b was not abrogated in presence of CVF is somewhat surprising (Fig. 5B) as this antibody did promote C1q recruitment on the surface of *S. aureus* (Fig. 5D). Another interesting finding is that mAb 8B9, which lacked any inhibitory activity toward ClfA-A ligands, is able to significantly promote the uptake of bacteria in whole blood, even in the absence of complement (Fig. 5AB).

**Fig. 5.**
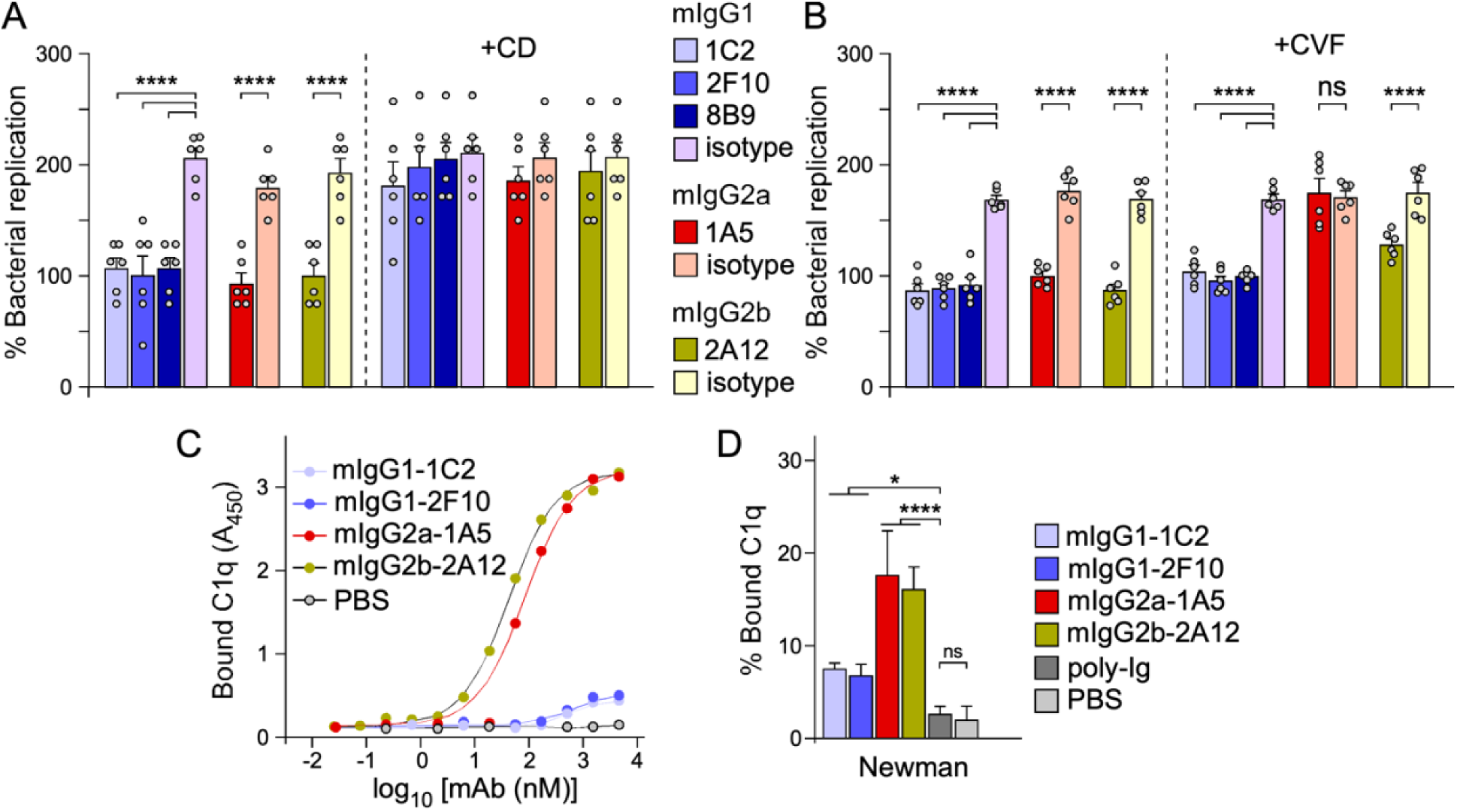
Opsonophagocytic and C1q binding activities of anti-ClfA antibodies. (A, B) **Staphylococcal survival in freshly drawn mouse blood** was assessed by inoculating mid-log phase bacteria (2.5 × 10^5^ CFU) into 0.5 ml of freshly drawn anti-coagulated mouse blood. At 0 min and 30 min, buffer was added to lyse host cells and release bacteria trapped in fibrin or NETs. Bacteria were counted by plating serial dilutions on solid medium and reported as % of viable counts with 100% representing bacterial counts at time 0. In panel A, one set of experiments was performed with blood pre-incubated with cytochalasin D (+ CD) to block phagocytic uptake. In panel B, one set of experiments was performed with blood pre-incubated with Cobra Venom Factor (+ CVF) to deplete complement. Data in each panel are from two independent experiments and are presented as mean ± SEM. Statistical significance was performed by Brown-Forsythe and Welch ANOVA followed by Dunnett’s T3 correction for multiple comparisons (****, *P* < 0.0001). Experiments were performed twice for reproducibility. (C, D) Association of mouse C1q with immobilized test antibodies (C) or immobilized Newman bacteria pre-incubated with test antibodies, poly-Ig or PBS (D). Bound C1q was detected using anti-mouse C1q-HRP and signals recorded as A_450_. In panel D, each A_450_ value (including PBS) was divided by the average absorbance value of the PBS control multiplied by 100 (A_450_ values were obtained in triplicates for each treatment); the data are presented as mean ± SEM and statistical significance calculated using One-way ANOVA with Tukey’s multiple comparison (****, *P* < 0.0001; *, *P* < 0.05; ns: not significant). Experiments in panels C and D were performed twice independently.

It has previously been shown that ClfA may also recruit complement factor I on the bacterial surface thereby increasing the cleavage of C3b into inactive C3b, an activity shown to correlate with reduced phagocytosis of staphylococci by purified human neutrophils (15, 63, 64). Complement factor I is also known as the C3b/C4b inactivator. Using ELISA, we confirmed that ClfA-A interacts with comnplement factor I (Table 2), however none of the mAbs was able to displace this interaction (Table 3).

### Anti-ClfA mAbs protect mice against blood stream challenges with *S. aureus*

A passive immunization experiment was performed to evaluate the protective activity of candidate mAbs in a mouse model of blood stream infection. Antibodies were administered 24 hours prior to intravenous challenge with a sub-lethal inoculum of *S. aureus.* In this model, bacteria that survive replication in blood disseminate to organ tissues to seed abscess lesions (11, 65, 66), a metastatic activity also observed following *S. aureus* bacteremia in humans (1, 2, 67). In infected mice, disease severity can be assessed by recording weight changes daily. As expected, significant weight losses were observed during the acute phase of disease (∼ first 6 days post-challenge) followed by a slow recovery (Fig. 6A-C).

**Fig. 6.**
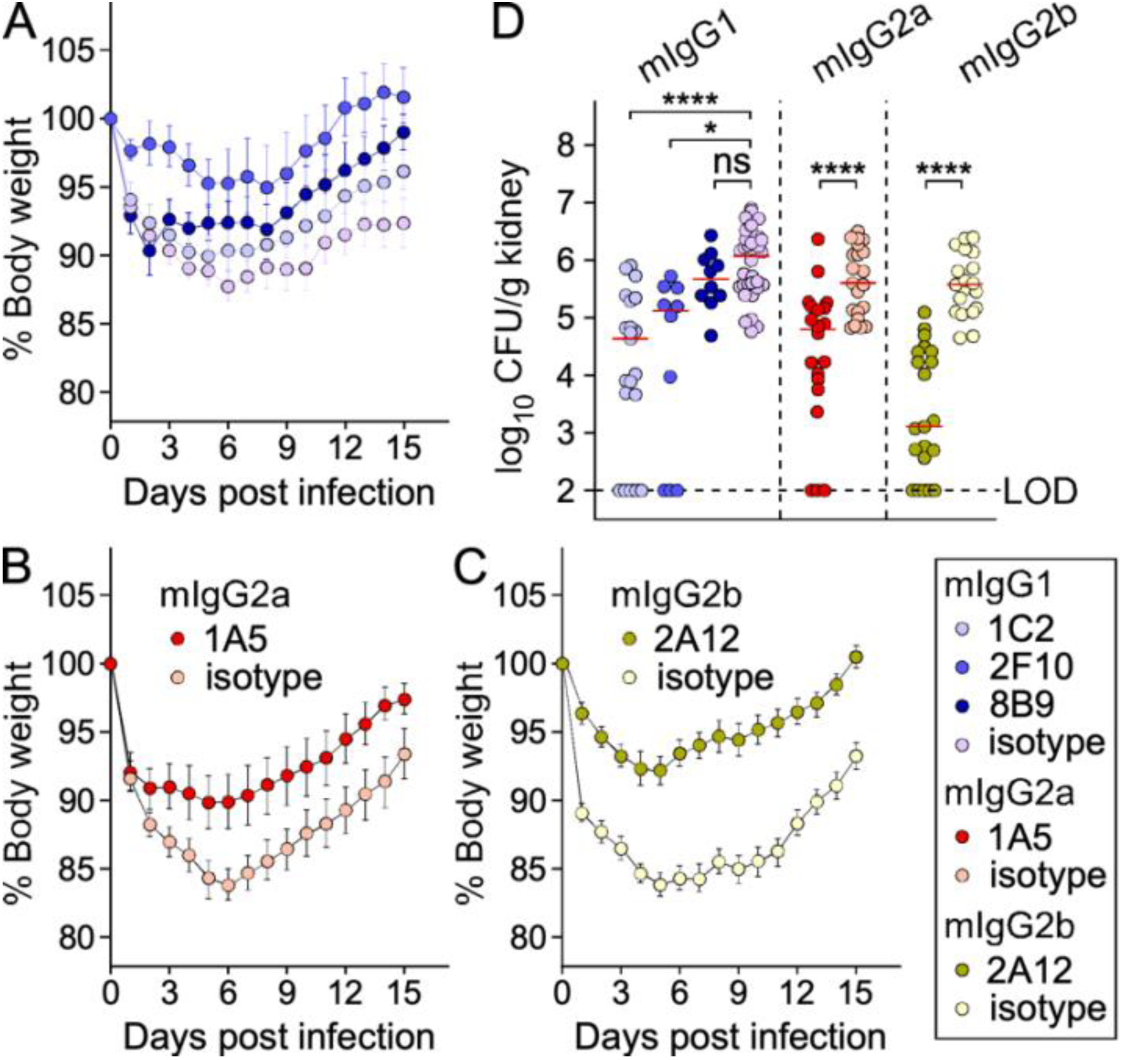
Protective activity of anti-ClfA mAbs following intravenous challenge of mice with *S. aureus.* Animals (*n*=8-12) were passively immunized with test or control antibodies (5 mg/kg IP) and challenged 24 hours later with a sub-lethal inoculum of strain USA300. Mice were weighed daily. (A-C) Weights were recorded daily, averaged, and changes reported as % with100% set on day 0 of the experiment (infection day). Data were split by isotypes for clarity. Statistical differences were calculated with the Kruskal-Wallis test followed by Dunn’s multiple test correction. (D) Bacterial loads in kidneys were enumerated by plating organs on day15 post infection. Bacterial counts are presented as median log_10_ CFU per g of kidney (LOD: limit of detection). Statistical significance was calculated using one-way ANOVA (mixed-model) with Gaussier-Greenhouse correction followed by Tukey’s multiple comparison test (****, *P* < 0.0001; *, *P* < 0.05; ns: not significant). A cumulative of at least two independent challenge studies are shown (testing of the 8B9 antibodies was only performed twice).

Animals were grouped by isotypes to better illustrate these changes. Of note, weight loss was less exacerbated with the passive administration of isotype control mIgG1 (Fig. 6A) as compared to mIgG2a (Fig. 6B) and mIgG2b (Fig. 6C) (Supplementary Table 3). As a result, alleviation of weight loss was more significant upon administration of 1A5-mIgG2a (Fig. 6B) or 2A12-mIgG2b (Fig. 6C) (Supplementary Table 3). Animals were killed on day 15 post-infection and bacterial burdens in kidneys (an organ where bacteria continue to replicate in deep-seated abscesses for weeks) were enumerated by plating serial dilutions of homogenized tissues. Animals pre-treated with 1C2-mIgG1, 2F10-mIgG1, 1A5-mIgG2a, 2A12-mIgG2b, showed significantly lower bacterial load in kidneys compared to isotype controls (Fig. 6D). Although, treatment with 2A12-mIgG2b resulted in the greatest reduction in the median bacterial loads, this reduction was no statistically significant as compared to treatment with the other three mAbs (Supplementary Table 3). Animals pre-treated with 8B9-mIgG showed no reduction in bacterial load in the kidneys of infected animals (Fig. 6D). Thus, although 8B9 displays opsonophagocytic activity in whole blood, this activity alone is not sufficient to reduce bacterial burdens in infected animals.

## DISCUSSION

*S. aureus* mutants lacking functional *clfA* display virulence defects in several mouse models of infection including septic arthritis, endocarditis, and abscess formation following blood stream dissemination of bacteria (11, 13, 19–22). These phenotypes have largely been correlated with the loss of staphylococcal binding to fibrinogen. Indeed, mice lacking the last five residues of the fibrinogen γ-chain (ClfA binding site) are more resistant to *S. aureus* sepsis (16, 68). Over twenty years ago, Patti and co-workers isolated mouse hybridomas MAb 12-9, MAb 35-052 and MAb 15EC6 using ClfA-A Newman as an immunogen (47). Only MAb 12-9, that bound ClfA on the surface of *S. aureus* and inhibited interaction with fibrinogen, conferred a modest survival benefit in BALB/c mice challenged intravenously with a lethal inoculum of strain Newman (47). Lack of protection by the other two antibodies was attributed to an inability to recognize ClfA on the surface of *S. aureus* (MAb 35-052) and to inhibit ClfA interaction with fibrinogen (MAb 15EC6), respectively (47). Here, we used the same antigen to isolate murine mAbs and exploited the N1, N2, N3 subdomains to further delinate the specificity of mAbs. Of five antibodies identified in this study, three recognized the N3 subdomain, one the N1 subdomain and one the full length ClfA-A domain but not the isolated N1, N2, N3 subdomains. We found that all the antibodies, with the exception of 8B9 (N1 specificity), were able to inhibit the interaction of ClfA with fibrinogen *in vitro.* However, the past twenty years of research have also revealed that ClfA binds to multiple ligands (12). In fact, it is now recognized that most MSCRAMMS have mutilple ligands, allowing the limited repertoire of surface proteins to carry out a vast array of host pathogen interactions (8, 9, 12). ClfA mediates interactions with fibrin, complement factor I and vWbp and vWF. In turn, these interactions promote bacterial agglutination, escape from phagocytes and complement, and bacterial adhesion. We found that with the exception of 8B9 (N1 specifcity), all the mAbs reduced bacterial agglutination and adhesion to host cells under flow, the latter inhibitory activity also correlating with the disruption of the ClfA-vWbp-vWF tether *in vitro*. However none of the mAbs could prevent the interaction between complement factor I and ClfA-A. Whether neutralizing such an interaction is important for protection during blood stream infection remains to be determined.

Patti and co-workers restricted their selection of mAbs to mIgG1 reasoning that this isotype is not captured by Staphylococcal protein A (SpA) on the surface of *S. aureus*. Indeed, SpA binds the Fc (fragment crystallizable) region of IgG with the hierarchy mIgG2a > mIgG2b > mIgG3 >>> mIgG1. SpA-mediated Fc binding diverts antibodies from their target and interferes with C1q recruitment, effectively preventing complement fixing and opsonophagocytosis of bacteria (69–71). mAbs examined in our study were of various subclasses allowing us to correlate bacterial killing in blood and in animals with the purported effector functions of antibodies. Indeed, mouse IgG Fc regions interact with various affinities with C1q and Fcγ receptors (FcγRs) that include activating, FcγRI, FcγRIII, FcγRIV, and inhibitory, FcγRIIb, receptors (72, 73). For example, while mIgG1 does not bind C1q, it can interact with the activating FcγRIII and inhibitory FcγRIIb receptors (72–74). Conversely, mIgG2a and mIgG2b trigger complement fixing on target cells and promote interactions with all activating FcγRs (72–74). When calculated as the A/I ratio representing the highest binding affinity for an activating (A) receptor over the binding affinity for the inhibitory (I) FcγRIIb, it ensues that mIgG2a has the highest A/I ratio, followed by mIgG2b, and mIgG1 (A/I values of 69, 7, and 0.1, respectivelty) (73). When interrogated for their ability to promote bacterial killing in whole blood, all the mAbs in our study were found to enhance bacterial uptake, including 8B9-mIgG1. Addition of CVF in this assay did not alter the activity of mIgG1 antibodies 1C2, 2F10, 8B9, but abrogated the activity of 1A5-mIgG2a. This is in agreement with the respective affinities of mIgG1 (negligible) and mIgG2a (high) toward C1q. Since the opsonophagocytic activity of 2A12-mIgG2b was not affected by CVF we surmise that in this assay, 2A12 may operate in an FcγR-dependent manner. All five antibodies were also tested in a passive immunization study whereby BALB/c mice were challenged intraveneously with *S. aureus* and killed 15 days post infection to enumerate bacteria in kidneys. This experiment revealed that treatment with all the antibodies, with the exception of 8B9-mIgG1, resulted in reduced bacterial burdens. Thus, it appears that the most effective mAbs have two activitities: they neutralize ClfA interactions with host ligands, and promote opsonophagocytic uptake of bacteria. In case of 2F10 and 1C2, it is also possible that some protection may have been contributed by the purported anti-inflammatory activity of the mIgG1 subclass (74, 75). Overall, results presented here echo our previous observations using 3F6, a mAb that neutralizes and binds SpA on the surface of *S. aureus.* We noted earlier that 3F6-mIgG1, 3F6-mIgG2a, 3F6-mIgG2b, and 3F6-mIgG3 provided similar reduction in bacterial replication in BALB/c whole blood as well as in animals infected for 15 days (76). This was not the case for experiments performed with C57BL/6J animals, antibody protection ranked as 3F6-mIgG2a > 3F6-mIgG1≥ 3F6-mIgG2b ≫ 3F6-mIgG3 (no protection) (76).

Other anti-ClfA mAbs described in the literature are human-based antibodies. For instance, mouse hybridoma MAb 12-9 was subsequently humanized using a human IgG1 backbone. These efforts yielded Tefibazumab (77). A phase II clinical trial with bacteremic patients compared the efficacy of Tefibazumab (Aurexis^TM^) and antibiotic treatment with placebo (24). However, composite clinical end point analysis did not detect differences between placebo and antibody (24). Subsequent structural studies revealed that the Fab region of Tefibazumab did not bind at the interface between N2 and N3 but rather at the head portion of N3, letting the authors of the study to conclude that ClfA has two fibrinogen-binding sites (78). The clinical failure of Tefibazumab was thus attributed to an incomplete ability to block fibrinogen binding. However, the ability of Tefibazumab to block interactions with other ligands such as fibrin, vWbp or complement factor I was also not tested. While it is possible that Tefibazumab did not meet correlates for ClfA neutralization, it is equally plausible that the constant region of the antibody may not have been optimal for activity in the human host. In another approach, Tkaczyk and colleagues isolated MAb 11H10 by immunizing VelocImmune mice with the N2-N3 subdomain of ClfA, *i.e.* the fibrinogen binding domain of ClfA (28). These genetically engineered mice produce human antibodies (79). Similarly to our anti-N3 antibodies, MAb 11H10 inhibited ClfA interactions with fibrinogen, reduced bacterial agglutination in plasma, and increased bacterial killing by HL-60 cells in the presence of human serum (we use anticoagulated whole blood). Yet, passive immunization of BALB/c mice resulted in a modest reduction in bacterial burdens in kidneys as compared to isotype control (∼0.5-log reduction 2 days post-blood stream challenge) (28). We observed a greater reduction in bacterial loads with 2A12-mIgG2b (∼2.5-log reduction 15 days post-challenge). While we cannot directly compare mouse and human mAbs, we reported earlier that the ability of humanized 3F6-hIgG1 to protect animals from *S. aureus* blood stream challenges and promote opsonophagocytic killing in human whole blood, was dependent on Asparagine 297 (Asn 297) glycosylation. Indeed, Asn297 in the constant region of hIgG1 is modified with a heptasaccharide core; variable additions of fucose, galactose, and sialic acid, modulate the effector functions of this antibody (80).

In conclusion, ClfA, a functionally well-characterized antigen of *S. aureus*, represents a great model to decipher antibody activities that correlate with protection against *S. aureus* in infected hosts. Whether non-neutralizing antibodies, such as those with N1-specificity, may exert diluting activity should be taken into consideration. Equally important is the appreciation that successful mAbs must neutralize most, if not all, ligand interactions promoted by ClfA as well as engage with Fc ligands for the effective clearance of bacteria.

## MATERIALS AND METHODS

### Ethics statement

The Institutional Animal Care and Use Committee at The University of Chicago reviewed, approved and supervised the protocols for animal experiments and the specific guidelines were followed during all procedures. All experiments involving *S. aureus* were performed in biosafety level 2 containment.

### Bacterial strains, hybridoma and mammalian cells, and growth media

Wild type *S. aureus* USA300, Newman and its mutants (*clfA, vwb*), were cultured in tryptic soy broth (TSB) or agar (TSA) at 37°C. *Escherichia coli* strains DH5α and BL21 harboring recombinant protein expression vector(s) were cultured in Luria-Bertani (LB) broth or agar with 100μg/ml carbenicillin at 37°C. Hybridoma cells were cultured in Iscove’s Modified Eagle’s Medium (IMDM) (Gibco) with 10% fetal bovine serum and 1X Pen Strep (Gibco) antibiotic solution additive. HUVECs were cultured in Endothelial cell ready-to-use growth medium (PromoCell) in a humidified incubator at 37°C and 5% CO_2_ injection. All bacterial mutant strains and hybridomas were from our laboratory collection.

### Production of mAbs against ClfA

mAbs against ClfA-A were generated as described previously (59, 81). Briefly, eight weeks old BALB/c mice were immunized with purified recombinant ClfA-A Newman (100 μg) diluted 1:1 in Freund’s adjuvant by intraperitoneal injection. Animals were boosted on days 21 and 42. Animals were bled on days 31 and 52 and screened for anti-ClfA antibodies. Mice with the highest immune-reactivity against ClfA were further boosted with 25 µg ClfA-A in PBS. Mice were harvested after three days and the isolated splenocytes were fused with the mouse myeloma cell line SP2/mIL-6. For selection, hybridomas were screened by ELISA for antigen-positive clones. For antibody production, hybridomas were cultured to a density of 10^6^ cells/ml in IMDM with 10% FBS until 90-95% confluence.

### Expression, and purification of recombinant proteins and mAbs

Recombinant A domain of ClfA protein (ClfA-A) from genomic DNA of *S. aureus* strains Newman, N315, WU1 and USA300 were amplified using primers ClfA-A-F:

CGCGCGGCAGCCATATGAGTGAAAATAGTGTTACGCAATCTGATAGC and ClfA-A-R: GTTAGCAGCCGGATCCCTTTTCGAACTGCGGGTGGCTCCACTCTGGAATTGGTTCAATTTCACC). ClfA-N1, N2 and N3 from Newman were amplified using the primer pairs N1-F: CGCGCGGCAGCCATATGAGTGAAAATAGTGTTACGCAATCTGATAGC, N1-R: GTTAGCAGCCGGATCCTGCCGCTAAACTAAATGCTCTCATTCTAGGC, N2-F: GTTAGCAGCCGGATCCTGCTTTTACATCATCTTTAGTATTTACATAGTCTGTAAATG, N2-R: CGCGCGGCAGCCATATGGTAGCTGCAGATGCACCG and N3-F: CGCGCGGCAGCCATATGACTTTGACCATGCCCGC and N3-R: GTTAGCAGCCGGATCCCTCTGGAATTGGTTCAATTTCACC respectively. The resulting gene fragments were cloned into pET15b vector using the *Nde*I *and BamH*I restriction sites in *E. coli* DH5α. To facilitate protein purification, six histidine residues (6-HIS) and Strep-tag II peptide (Strep) consisting of Trp-Ser-His-Pro-Gln-Phe-Glu-Lys were engineered at the N- and C-termini of each gene product (some variants carried both or either one of the tags for purification and ELISA detection). The resulting plasmids were sequenced and transferred to *E. coli* BL21. The resulting strains were propagated in liquid cultures to mid-logarithmic growth before addition of 1mM Isopropyl β-D-1-thiogalactopyranoside (IPTG) to induce the production of recombinant proteins. Cleared cell lysates were passed over Ni-NTA resin followed by Streptactin resin following manufacturer’s instructions. All antibodies were purified from the supernatants of hybridomas cell cultures grown in serum-free Freestyle 293 medium by affinity chromatography over Protein G sepharose (Cytiva) following the manufacturer’s protocol and dialyzed against PBS (pH 7.2) (76). Hybridoma cells producing 2F10-mIgG1 were purified over ClfA-A crosslinked to AminoLink^TM^ Plus coupling resin (Thermofisher). Proteins and antibodies were dialized against PBS and kept at 2 mg/ml for subsequent downstream *in vitro* and *in vivo* assays.

### Enzyme-linked immunosorbent assays (ELISAs)

ELISA-based assays were used to measure interactions between mAbs and ClfA-A antigen, ClfA-A and ligands as well as to assess the ability of antibodies to block interactions (competition assays). Briefly, to measure binding interactions between two partners, Nunc Maxisorp 96-well plates (Thermofisher) were coated with either 100 ng recombinant antigens (ClfA-A, N1, N2 or N3 variants), 100 ng mAbs, 100 ng human vWF (Sigma), or bacterial cells at 1 × 10^6^ CFU (per well) in 100 mM carbonate/bi-carbonate buffer (pH 9.6) at 4 °C overnight (for proteins) and 37°C for 2 h (for bacterial cells). After blocking the wells with 1% (w/v) bovine serum albumin (BSA) in phosphate buffer containing tween-20 (PBST) for 1 h, 100 μl serial dilutions of either test antibodies (20 μg/ml), human fibrinogen (4 mg/ml; Sigma), human complement factor I (20 μg/ml; MiliporeSigma), recombinant ClfA-A proteins (20 μg/ml), or recombinant vWbp (5 μg/ml) were added to the wells. To evaluate C1q binding to mAbs, mouse C1q (20 μg/ml; CompTech, M099) was added to the plates coated with mAbs. To assess the inhibitory activity of antibodies in blocking interactions (competition experiments), 50 μl of serial dilutions of 20 μg/ml test antibodies were mixed with 50 μl of either human fibrinogen (4 mg/ml), human complement factor I (20 μg/ml), vWbp (5 μg/ml) or vWbp plus human vWF, and then added to plates coated with ClfA-A. Bound antibodies or fibrinogen were determined by incubating with HRP conjugated anti-mouse IgG antibody (Southern Biotech; 1:10,000 dilution) and HRP conjugated anti-human fibrinogen antibody (Rockland, 1:5000), respectively. Bound recombinant ClfA-A and N subdomains were detected with HRP conjugated anti-StrepII antibody (Genscript; 1:5000 dilution) or anti-His-HRP antibody (Abcam; 1:10,1000 dilution). Bound mouse C1q was detected with anti-mouse C1q antibody JL-1 (Invitrogen) that had been conjugated using LYNX rapid antibody HRP conjugation kit (BioRad; 1:50 dilution). Bound complement factor I was detected using rabbit anti-factor I polyclonal antibody (Invitrogen, 1:5000 dilution) and goat anti-rabbit HRP antibody (Southern Biotech, 1:5000). For vWF-vWbp-ClfA-A interaction, recombinant ClfA-A carrying only His tag was added to the reaction and bound vWbp was assessed with anti-StrepII antibody (Genscript; 1:5000 dilution). To determine if mAbs compete for the same epitope, serial dilutions of unconjugated 1A5, 1C2 and 2A12 were added as a competitor to HRP-conjugated 2A12 (using LYNX rapid antibody HRP conjugation kit, BioRad) in 5:1 ratio to ELISA plates coated with isolated N3. To evaluate the presence of antibodies against ClfA in human sera, serial dilutions of IVIG (Privigen® 10% Liquid Intravenous Immunoglobulin, CSL Behring) were prepared by an initial dilution of 1:10,000 in blocking buffer, followed by seven 1:5 serial dilutions. One hundred microliters of each dilution was added to wells previously coated with 100 ng recombinant ClfA-A or N1, N2, or N3 variants. Binding was detected by incubating with HRP conjugated anti-human IgG (Promega, 1:10,000) for 1 h at room temperature. All plates were developed using OptEIA reagent (BD Biosciences) and signals quantified using a plate reader by recording absorbances at 450 nm (*A*_450_). Half maximum titers defined as the reciprocal of the dilution yielding 50% of the maximal absorbance, were determined using GraphPad Prism 10.

### *In vitro* agglutination assay

The bacterial agglutination assay in plasma was performed as described previously (13, 82). In brief, 10 ml of *S. aureus* Newman grown for 16 hours was centrifuged, followed by wash with PBS and resuspended in PBS (normalized to absorbance at 600 nm *A*_600_=4). Bacterial cells were stained with SYTO 9 (1:500) (Invitrogen) for 15 min away from light. Then the cells were resuspended in 1ml PBS after two washes and test antibodies added at a final concentration of 10 μg/ml. These bacterial suspensions were mixed 1:1 volumetrically with anticoagulated mouse plasma on a glass slide followed by 30 min incubation at room temperature. Samples were observed under an inverted fluorescence microscope (Lyca) and 10 different fields of view were captured for quantification of agglutinated area using QPath software.

### Adhesion assay

The adhesion of *S. aureus* to host cells was quantified by an *in vitro* perfusion assay adapted from (83). Briefly, human umbilical vein endothelial cells (HUVECs) were expanded in a culture flask and grown to 80% confluency. 50μl of the cells were transferred into the channels of μSlide VI 0.4 (Ibidi). The cells were allowed to adhere overnight followed by addition of Ca+ ionophore (Sigma) to induce human von Willebrand Factor (vWF) secretion [10]. *S. aureus* Newman cultures were grown to mid-log phase. Cells were washed, resuspended in PBS and stained with SYTO 9 (1:500), washed again and resuspended in PBS at *A*_600_ =0.42 to yield ∼1×10^8^ cells per ml suspension. Test antibodies were added at 10 μg/ml to the bacterial suspension. The resulting mixture of bacteria and antibodies were flowed at 1ml/min through the micro slide channels containing HUVECs for 15 min using a peristaltic pump. The micro slide channels were subsequently washed with PBS at 1ml/min for 15 min and allowed to dry. Bound staphylococcal cells were visualized under an inverted fluorescence microscope (Lyca) and 10 different fields of view were captured. The number of cells were counted and represented as percentage of bound cells using Prism 10 (GraphPad).

### Bacterial survival in mouse whole blood

The replication of bacteria in freshly drawn blood in the presence of test antibodies was measured as described previously (59, 84). Briefly, 10μg/ml test antibodies were added to 500 μl of freshly drawn BALB/cJ mouse blood anticoagulated with 10 μg/ml heparin. 50 μl of a bacterial suspension (2.5×10^5^ CFU/ml) was added. When noted, the blood was preincubated with Cytochalasin D (CD) or Cobra Venom Factor (CVF) for 10 min. Otherwise, tubes inoculated with bacteria were incubated at 37 °C for 30 min. Next, 500 μl SK Buffer containing 0.5% saponin, 100 U streptokinase, 50 μg trypsin, 1 μg DNase, and 5 μg RNase in PBS, was added to each sample followed by incubation at 37°C for 10 mins to release live bacteria from agglutinates, NETs or intracellular compartments. Sample aliquots were plated on agar for CFU enumeration in triplicates and all experiments were repeated at least twice independently.

### Passive immunization studies

BALB/cJ mice (6-8 weeks old; 50% female, 50% male) were immunized with test antibodies (5 mg/kg) via intra-peritoneal injection. Mice were anesthetized with isoflurane and sub-lethal 5×10^6^ CFU of *S. aureus* were retro-orbitally injected 24-hour post immunization in groups of 8 to 12 animals. The health of the mice was monitored for signs of acute disease and by recording weight daily. For CFU enumeration of bacteria in kidneys, animals were killed at day 15 post sub-lethal challenge. All challenge experiments were repeated at least twice.

### Statistical analyses

Data were analyzed for statistical significance at 95% confidence interval using one-way ANOVA with Tukey’s multiple comparison test between samples and the controls with the exception of the study examining vWF and vWbp interactions with ClfA-A for which the analysis was performed with the unpaired *t* test (Table 2). Binding assays and competition assays using ELISAs were plotted and the curve was fitted with non-linear (least squares) regression. Differences in agglutination areas and bacterial replication in whole blood were analyzed using Brown-Forsythe and Welch ANOVA followed by Dunnett’s T3 correction for multiple comparisons. Changes in body weight were reported as the average of the body weights of mice across treatment groups and analyzed using Kruskal-Wallis test followed by Dunn’s multiple test correction for statistical significance. Bacterial burdens in kidneys of infected mice were compared using one-way ANOVA (mixed-model) with Gaussier-Greenhouse correction followed by Tukey’s multiple comparison test. All data were analyzed with Prism 10 (GraphPad Software, Inc.), and *p* values less than 0.05 were considered significant and marked with asterisks on the graphs.

## Acknowledgements

We thank Andrea Dedent, Xinhai Chen, Hwan Keun Kim, Vilasack Thammavongsa, Lena Thomer for their support and insights on this work, Derek Elli, Haley Gula and Anastasia Tomatsidou for facilitating experimental work. This research was supported by grants AI110937 and AI1485437 from the National Institute of Allergy and Infectious Diseases, Infectious Disease Branch.

## Supplementary Figures

**Fig. S1.**
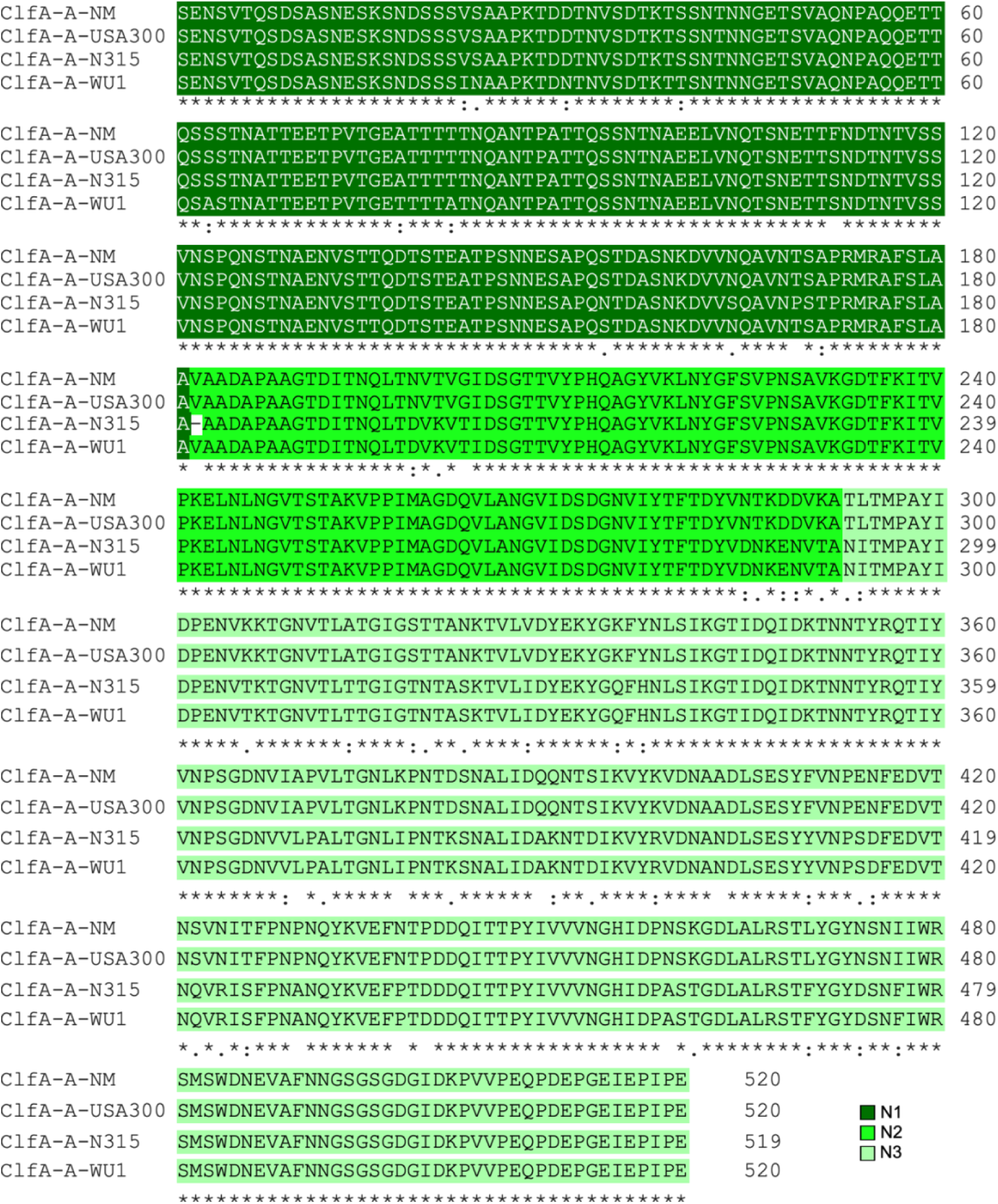
Sequence alignments of ClfA-A proteins. Sequence alignments of ClfA-A domains from strains Newman (NM), USA300, N315 and WU1 were generated using Clustal Omega (https://www.ebi.ac.uk/jdispatcher/msa/clustalo). The N1, N2 and N3 subdomains are shown in dark, bright and light green, respectively. The conservation score symbols shown below each position in the alignment indicate residues that are: (*),identical; (:), conserved; (.), semi-conserved. The absence of score symbol (blank) indicates non-conserved residues.

**Fig. S2.**
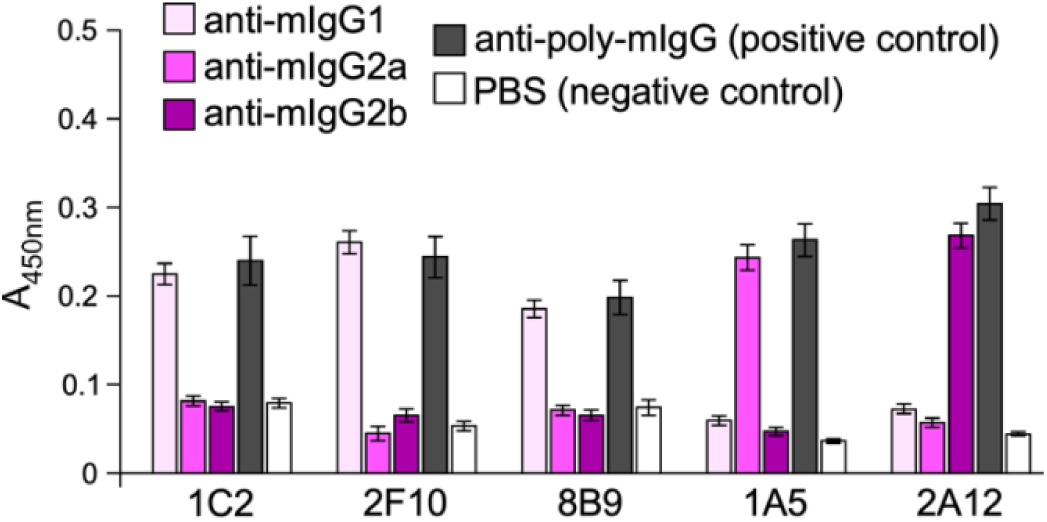
Isotype determination of experimental mAbs. Candidate antibodies, 1C2, 2F10, 8B9, 1A5 and 2A12 bound to 96-well plates were detected with biotin-tagged anti-mIgG1, 2a, 2b and poly-mIgG, followed by streptavidin-HRP conjugate secondary antibody.

**Fig. S3.**
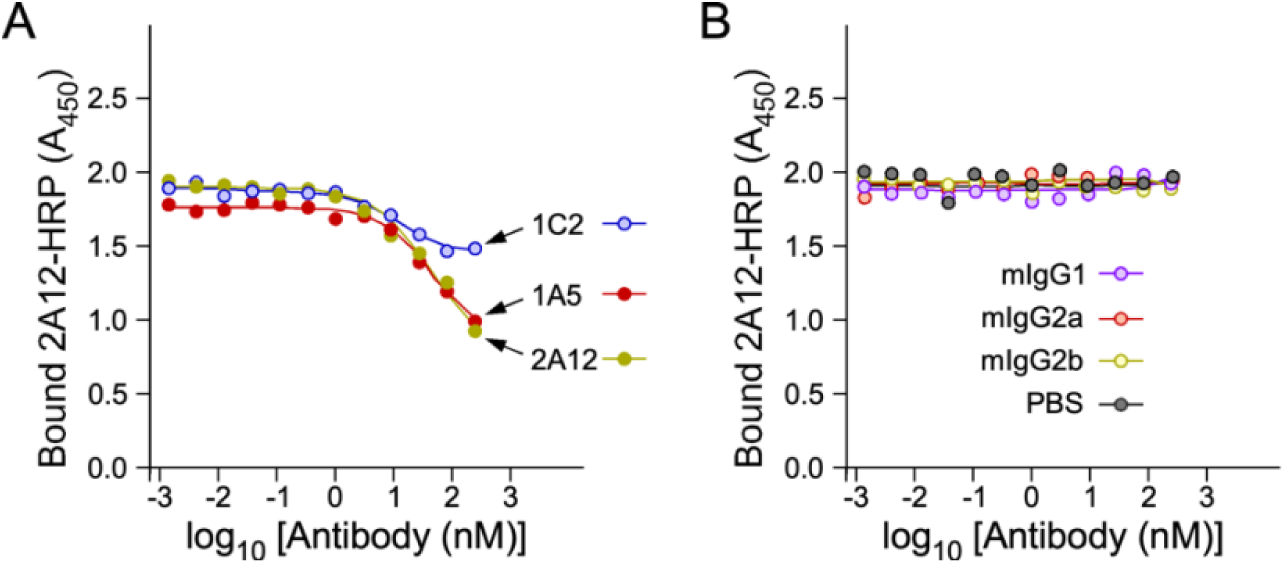
Competive ELISA experiment between N3-specific mAbs. HRP labeled 2A12 (2A12-HRP) was mixed with serially diluted unlabeled 1A5, 1C2 and 2A12 (**A**) or isotype control antibodies mIgG1, 2a, 2b or PBS (**B**) and added to ELISA plates coated with the N3 subdomain. Bound 2A12-HRP was assessed by recording absorbance at 450 nm (*A*_450_). A representative of three experiments is shown.

## REFERENCES

1. Lowy FD. 1998. *Staphylococcus aureus* infections. New Engl J Med 339:520–532.

2. Tong SY, Davis JS, Eichenberger E, Holland TL, Fowler VG, Jr. 2015. *Staphylococcus aureus* infections: epidemiology, pathophysiology, clinical manifestations, and management. Clin Microbiol Rev 28:603–61.

3. Fitzgerald JR. 2012. Livestock-associated *Staphylococcus aureus*: origin, evolution and public health threat. Trends Microbiol 20:192–198.

4. Haag AF, Fitzgerald JR, Penades JR. 2019. Staphylococcus aureus in Animals. Microbiol Spectr 7.

5. Howden BP, Giulieri SG, Wong Fok Lung T, Baines SL, Sharkey LK, Lee JYH, Hachani A, Monk IR, Stinear TP. 2023. Staphylococcus aureus host interactions and adaptation. Nat Rev Microbiol 21:380–395.

6. Foster TJ, Hook M. 1998. Surface protein adhesins of Staphylococcus aureus. Trends Microbiol 6:484–8.

7. Foster TJ, Geoghegan JA, Ganesh VK, Hook M. 2014. Adhesion, invasion and evasion: the many functions of the surface proteins of Staphylococcus aureus. Nat Rev Microbiol 12:49–62.

8. Marraffini LA, DeDent AC, Schneewind O. 2006. Sortases and the art of anchoring proteins to the envelopes of gram-positive bacteria. Microbiol Mol Biol Rev 70:192–221.

9. Schneewind O, Missiakas D. 2019. Sortases, Surface Proteins, and Their Roles in Staphylococcus aureus Disease and Vaccine Development. Microbiol Spectr 7.

10. Mazmanian SK, Liu G, Jensen ER, Lenoy E, Schneewind O. 2000. *Staphylococcus aureus* sortase mutants defective in the display of surface proteins and in the pathogenesis of animal infections. Proc Natl Acad Sci USA 97:5510–5515.

11. Cheng AG, Kim HK, Burts ML, Krausz T, Schneewind O, Missiakas DM. 2009. Genetic requirements for *Staphylococcus aureus* abscess formation and persistence in host tissues. FASEB J 23:3393–3404.

12. Foster TJ. 2019. The MSCRAMM Family of Cell-Wall-Anchored Surface Proteins of Gram-Positive Cocci. Trends Microbiol 27:927–941.

13. McAdow M, Kim HK, DeDenta AC, Hendrickx APA, Schneewind O, Missiakas DM. 2011. Preventing *Staphylococcus aureus* sepsis through the inhibition of its agglutination in blood. PLoS Pathog 7:e1002307.

14. Palmqvist N, Patti JM, Tarkowski A, Josefsson E. 2004. Expression of staphylococcal clumping factor A impedes macrophage phagocytosis. Microb Infect 6:188–195.

15. Hair PS, Echague CG, Sholl AM, Watkins JA, Geoghegan JA, Foster TJ, Cunnion KM. 2010. Clumping factor A interaction with complement factor I increases C3b cleavage on the bacterial surface of *Staphylococcus aureus* and decreases complement-mediated phagocytosis. Infect Immun 78:1717–1727.

16. Flick MJ, Du X, Prasad JM, Raghu H, Palumbo JS, Smeds E, Höök M, Degen JL. 2013. Genetic elimination of the binding motif on fibrinogen for the *S. aureus* virulence factor ClfA improves host survival in septicemia. Blood in press.

17. Claes J, Liesenborghs L, Peetermans M, Veloso TR, Missiakas D, Schneewind O, Mancini S, Entenza JM, Hoylaerts MF, Heying R, Verhamme P, Vanassche T. 2017. Clumping factor A, von Willebrand factor-binding protein and von Willebrand factor anchor Staphylococcus aureus to the vessel wall. J Thromb Haemost 15:1009–1019.

18. Claes J, Ditkowski B, Liesenborghs L, Veloso TR, Entenza JM, Moreillon P, Vanassche T, Verhamme P, Hoylaerts MF, Heying R. 2018. Assessment of the Dual Role of Clumping Factor A in S. Aureus Adhesion to Endothelium in Absence and Presence of Plasma. Thromb Haemost 118:1230–1241.

19. Josefsson E, Hartford O, O’Brien L, Patti JM, Foster TJ. 2001. Protection against experimental *Staphylococcus aureus* arthritis by vaccination with clumping factor A, a novel virulence determinant. J Infect Dis 184:1572–1580.

20. Arrecubieta C, Asai T, Bayern M, Loughman A, Fitzgerald JR, Shelton CE, Baron HM, Dang NC, Deng MC, Naka Y, Foster TJ, Lowy FD. 2006. The role of Staphylococcus aureus adhesins in the pathogenesis of ventricular assist device-related infections. J Infect Dis 193:1109–19.

21. Moreillon P, Entenza JM, Francioli P, McDevitt D, Foster TJ, François P, Vaudaux P. 1995. Role of *Staphylococcus aureus* coagulase and clumping factor in pathogenesis of experimental endocarditis. Infect Immun 63:4738–4743.

22. Liesenborghs L, Meyers S, Lox M, Criel M, Claes J, Peetermans M, Trenson S, Vande Velde G, Vanden Berghe P, Baatsen P, Missiakas D, Schneewind O, Peetermans WE, Hoylaerts MF, Vanassche T, Verhamme P. 2019. Staphylococcus aureus endocarditis: distinct mechanisms of bacterial adhesion to damaged and inflamed heart valves. Eur Heart J 40:3248–3259.

23. Nour El-Din AN, Shkreta L, Talbot BG, Diarra MS, Lacasse P. 2006. DNA immunization of dairy cows with the clumping factor A of Staphylococcus aureus. Vaccine 24:1997–2006.

24. Weems JJ, Jr., Steinberg JP, Filler S, Baddley JW, Corey GR, Sampathkumar P, Winston L, John JF, Kubin CJ, Talwani R, Moore T, Patti JM, Hetherington S, Texter M, Wenzel E, Kelley VA, Fowler VG, Jr. 2006. Phase II, randomized, double-blind, multicenter study comparing the safety and pharmacokinetics of tefibazumab to placebo for treatment of Staphylococcus aureus bacteremia. Antimicrob Agents Chemother 50:2751–5.

25. John JF, Jr. 2006. Drug evaluation: tefibazumab--a monoclonal antibody against staphylococcal infection. Curr Opin Mol Ther 8:455–60.

26. Hawkins J, Kodali S, Matsuka YV, McNeil LK, Mininni T, Scully IL, Vernachio JH, Severina E, Girgenti D, Jansen KU, Anderson AS, Donald RG. 2012. A recombinant clumping factor A-containing vaccine induces functional antibodies to *Staphylococcus aureus* that are not observed after natural exposure. Clin Vaccine Immunol 19:1641–1650.

27. Scully IL, Timofeyeva Y, Keeney D, Matsuka YV, Severina E, McNeil LK, Nanra J, Hu G, Liberator PA, Jansen KU, Anderson AS. 2015. Demonstration of the preclinical correlate of protection for *Staphylococcus aureus* clumping factor A in a murine model of infection. Vaccine 33:5452–5457.

28. Tkaczyk C, Hamilton MM, Sadowska A, Shi Y, Chang CS, Chowdhury P, Buonapane R, Xiao X, Warrener P, Mediavilla J, Kreiswirth B, Suzich J, Stover CK, Sellman BR. 2016. Targeting Alpha Toxin and ClfA with a Multimechanistic Monoclonal-Antibody-Based Approach for Prophylaxis of Serious Staphylococcus aureus Disease. MBio 7.

29. Tkaczyk C, Kasturirangan S, Minola A, Jones-Nelson O, Gunter V, Shi YY, Rosenthal K, Aleti V, Semenova E, Warrener P, Tabor D, Stover CK, Corti D, Rainey G, Sellman BR. 2017. Multimechanistic Monoclonal Antibodies (MAbs) Targeting Staphylococcus aureus Alpha-Toxin and Clumping Factor A: Activity and Efficacy Comparisons of a MAb Combination and an Engineered Bispecific Antibody Approach. Antimicrob Agents Chemother 61.

30. Alabdullah HA, Overgaard E, Scarbrough D, Williams JE, Mohammad Mousa O, Dunn G, Bond L, McGuire MA, Tinker JK. 2020. Evaluation of the Efficacy of a Cholera-Toxin-Based Staphylococcus aureus Vaccine against Bovine Intramammary Challenge. Vaccines (Basel) 9.

31. Scully IL, Timofeyeva Y, Illenberger A, Lu P, Liberator PA, Jansen KU, Anderson AS. 2021. Performance of a Four-Antigen Staphylococcus aureus Vaccine in Preclinical Models of Invasive S. aureus Disease. Microorganisms 9.

32. Nguyen NTQ, Doan TNM, Sato K, Tkaczyk C, Sellman BR, Diep BA. 2023. Monoclonal antibodies neutralizing alpha-hemolysin, bicomponent leukocidins, and clumping factor A protected against Staphylococcus aureus-induced acute circulatory failure in a mechanically ventilated rabbit model of hyperdynamic septic shock. Front Immunol 14:1260627.

33. Ponnuraj K, Bowden MG, Davis S, Gurusiddappa S, Moore D, Choe D, Xu Y, Hook M, Narayana SV. 2003. A “dock, lock, and latch” structural model for a staphylococcal adhesin binding to fibrinogen. Cell 115:217–228.

34. Bowden MG, Heuck AP, Ponnuraj K, Kolosova E, Choe D, Gurusiddappa S, Narayana SV, Johnson AE, Höök M. 2008. Evidence for the “dock, lock, and latch” ligand binding mechanism of the staphylococcal microbial surface component recognizing adhesive matrix molecules (MSCRAMM) SdrG. J Biol Chem 283:638–647.

35. Deivanayagam CC, Wann ER, Chen W, Carson M, Rajashankar KR, Höök M, Narayana SV. 2002. A novel variant of the immunoglobulin fold in surface adhesins of *Staphylococcus aureus*: crystal structure of the fibrinogen-binding MSCRAMM, clumping factor A. EMBO J 21:6660–6672.

36. McDevitt D, Nanavaty T, House-Pompeo K, Bell E, Turner N, McIntire L, Foster T, Höök M. 1997. Characterization of the interaction between the *Staphylococcus aureus* clumping factor (ClfA) and fibrinogen. Eur J Biochem 247:416–424.

37. Ganesh VK, Rivera JJ, Smeds E, Ko Y-P, Bowden MG, Wann ER, Gurusidappa S, Fitzgerald JR, Höök M. 2008. A structural model of the *Staphylococcus aureus* ClfA-fibrinogen interaction opens new avenues for the design of anti-staphylococcal therapeutics. PLoS Pathog 4:e1000226.

38. Doolittle RF. 2003. Structural basis of the fibrinogen-fibrin transformation: contributions from X-ray crystallography. Blood Rev 17:33–41.

39. McDevitt D, Francois P, Vaudaux P, Foster TJ. 1994. Molecular characterization of the clumping factor (fibrinogen receptor) of *Staphylococcus aureus*. Mol Microbiol 11:237–248.

40. Ashraf S, Cheng J, Zhao X. 2017. Clumping factor A of Staphylococcus aureus interacts with AnnexinA2 on mammary epithelial cells. Sci Rep 7:40608.

41. McAdow M, Missiakas DM, Schneewind O. 2012. *Staphylococcus aureus* secretes coagulase and von Willebrand factor binding protein to modify the coagulation cascade and establish host infections. J Innate Immun 4:141–148.

42. Baba T, Bae T, Schneewind O, Takeuchi F, Hiramatsu K. 2008. Genome sequence of Staphylococcus aureus strain Newman and comparative analysis of staphylococcal genomes: polymorphism and evolution of two major pathogenicity islands. J Bacteriol 190:300–10.

43. Hazenbos WL, Kajihara KK, Vandlen R, Morisaki JH, Lehar SM, Kwakkenbos MJ, Beaumont T, Bakker AQ, Phung Q, Swem LR, Ramakrishnan S, Kim J, Xu M, Shah IM, Diep BA, Sai T, Sebrell A, Khalfin Y, Oh A, Koth C, Lin SJ, Lee BC, Strandh M, Koefoed K, Andersen PS, Spits H, Brown EJ, Tan MW, Mariathasan S. 2013. Novel staphylococcal glycosyltransferases SdgA and SdgB mediate immunogenicity and protection of virulence-associated cell wall proteins. PLoS Pathog 9:e1003653.

44. Thomer L, Becker S, Emolo C, Quach A, Kim HK, Rauch S, Anderson M, Leblanc JF, Schneewind O, Faull KF, Missiakas D. 2014. N-acetylglucosaminylation of serine-aspartate repeat proteins promotes *Staphylococcus aureus* bloodstream infection. J Biol Chem 289:3478–86.

45. Murphy E, Lin SL, Nunez L, Andrew L, Fink PS, Dilts DA, Hoiseth SK, Jansen KU, Anderson AS. 2011. Challenges for the evaluation of Staphylococcus aureus protein based vaccines: monitoring antigenic diversity. Hum Vaccin 7 Suppl:51–9.

46. Brady RA, Mocca CP, Burns DL. 2013. Immunogenicity analysis of Staphylococcus aureus clumping factor A genetic variants. Clin Vaccine Immunol 20:1338–40.

47. Hall AE, Domanski PJ, Patel PR, Vernachio JH, Syribeys PJ, Gorovits EL, Johnson MA, Ross JM, Hutchins JT, Patti JM. 2003. Characterization of a protective monoclonal antibody recognizing *Staphylococcus aureus* MSCRAMM protein clumping factor A. Infect Immun 71:6864–6870.

48. Diep BA, Gill SR, Chang RF, Phan TH, Chen JH, Davidson MG, Lin F, Lin J, Carleton HA, Mongodin EF, Sensabaugh GF, Perdreau-Remington F. 2006. Complete genome sequence of USA300, an epidemic clone of community-acquired meticillin-resistant *Staphylococcus aureus*. Lancet 367:731–739.

49. Kuroda M, Ohta T, Uchiyama I, Baba T, Yuzawa H, Kobayashi I, Cui L, Oguchi A, Aoki K, Nagai Y, Lian J, Ito T, Kanamori M, Matsumaru H, Maruyama A, Murakami H, Hosoyama A, Mitsutani-Ui Y, Kobayashi N, Sawano T, Inoue R, Kaito C, Sekimizu K, Hirakawa H, Kuhara S, Goto S, Yabuzaki J, Kanehisa M, Yamashita A, Oshima K, Furuya K, Yoshino C, Shiba T, Hattori M, Ogasawara N, Hayashi H, Hiramatsu K. 2001. Whole genome sequencing of meticillin-resistant *Staphylococcus aureus*. Lancet 357:1225–1240.

50. Sun Y, Emolo C, Holtfreter S, Wiles S, Kreiswirth B, Missiakas D, Schneewind O. 2018. Staphylococcal protein A contributes to persistent colonization of mice with Staphylococcus aureus. J Bacteriol doi:10.1128/JB.00735-17.

51. Cullum E, Perez-Betancourt Y, Shi M, Gkika E, Schneewind O, Missiakas D, Golovkina T. 2024. Deficiency in non-classical major histocompatibility class II-like molecule, H2-O confers protection against Staphylococcus aureus in mice. PLoS Pathog 20:e1012306.

52. Birch-Hirschfeld L. 1934. Über die Agglutination von Staphylokokken durch Bestandteile des Säugetierblutplasmas. Klinische Woschenschrift 13:331.

53. Cheng AG, McAdow M, Kim HK, Bae T, Missiakas DM, Schneewind O. 2010. Contribution of coagulases towards *Staphylococcus aureus* disease and protective immunity. PLoS Pathog 6:e1001036.

54. McDonnell CJ, Garciarena CD, Watkin RL, McHale TM, McLoughlin A, Claes J, Verhamme P, Cummins PM, Kerrigan SW. 2016. Inhibition of major integrin alpha(V) beta(3) reduces Staphylococcus aureus attachment to sheared human endothelial cells. J Thromb Haemost 14:2536–2547.

55. Claes J, Vanassche T, Peetermans M, Liesenborghs L, Vandenbriele C, Vanhoorelbeke K, Missiakas D, Schneewind O, Hoylaerts MF, Heying R, Verhamme P. 2014. Adhesion of *Staphylococcus aureus* to the vessel wall under flow is mediated by von Willebrand factor-binding protein. Blood 124:1669–76.

56. Huang J, Roth R, Heuser JE, Sadler JE. 2009. Integrin alpha(v)beta(3) on human endothelial cells binds von Willebrand factor strings under fluid shear stress. Blood 113:1589–97.

57. Viljoen A, Viela F, Mathelie-Guinlet M, Missiakas D, Pietrocola G, Speziale P, Dufrene YF. 2021. Staphylococcus aureus vWF-binding protein triggers a strong interaction between clumping factor A and host vWF. Commun Biol 4:453.

58. Bjerketorp J, Nilsson M, Ljungh A, Flock JI, Jacobsson K, Frykberg L. 2002. A novel von Willebrand factor binding protein expressed by *Staphylococcus aureus*. Microbiology 148:2037–2044.

59. Thomer L, Emolo C, Thammavongsa V, Kim HK, McAdow ME, Yu W, Kieffer M, Schneewind O, Missiakas D. 2016. Antibodies against a secreted product of *Staphylococcus aureus* trigger phagocytic killing. J Exp Med 213:293–301.

60. Vogel C-W, Fritzinger DC. 2010. Cobra venom factor: structure, function, and humanization for therapeutic complement depletion. Toxicon 56:1198–1222.

61. Leatherbarrow RJ, Dwek RA. 1984. Binding of complement subcomponent C1q to mouse IgG1, IgG2a and IgG2b: a novel C1q binding assay. Mol Immunol 21:321–7.

62. Chemouny JM, Hurtado-Nedelec M, Flament H, Ben Mkaddem S, Daugas E, Vrtovsnik F, Berthelot L, Monteiro RC. 2016. Protective role of mouse IgG1 in cryoglobulinaemia; insights from an animal model and relevance to human pathology. Nephrology Dialysis Transplantation 31:1235–1242.

63. Cunnion KM, Buescher ES, Hair PS. 2005. Serum complement factor I decreases *Staphylococcus aureus* phagocytosis. J Lab Clin Med 146:279–286.

64. Hair PS, Ward MD, Semmes OJ, Foster TJ, Cunnion KM. 2008. Staphylococcus aureus clumping factor A binds to complement regulator factor I and increases factor I cleavage of C3b. J Infect Dis 198:125–33.

65. Cheng AG, DeDent AC, Schneewind O, Missiakas D. 2011. A play in four acts: Staphylococcus aureus abscess formation. Trends Microbiol 19:225–32.

66. Thomer L, Schneewind O, Missiakas D. 2016. Pathogenesis of *Staphylococcus aureus* Bloodstream Infections. Annu Rev Pathol 11:343–64.

67. Corey GR. 2009. Staphylococcus aureus bloodstream infections: definitions and treatment. Clin Infect Dis 48 Suppl 4:S254–9.

68. Negron O, Weggeman M, Grimbergen J, Clark EG, Abrahams S, Hur WS, Koopman J, Flick MJ. 2023. Fibrinogen gamma’ promotes host survival during Staphylococcus aureus septicemia in mice. J Thromb Haemost 21:2277–2290.

69. Forsgren A, Quie PG. 1974. Effects of staphylococcal protein A on heat labile opsonins. J Immunol 112:1177–1180.

70. Falugi F, Kim HK, Missiakas DM, Schneewind O. 2013. The role of protein A in the evasion of host adaptive immune responses by *Staphylococcus aureus* mBio 4:e00575–13.

71. Cruz AR, Boer MAD, Strasser J, Zwarthoff SA, Beurskens FJ, de Haas CJC, Aerts PC, Wang G, de Jong RN, Bagnoli F, van Strijp JAG, van Kessel KPM, Schuurman J, Preiner J, Heck AJR, Rooijakkers SHM. 2021. Staphylococcal protein A inhibits complement activation by interfering with IgG hexamer formation. Proc Natl Acad Sci U S A 118.

72. Neuberger MS, Rajewsky K. 1981. Activation of mouse complement by monoclonal mouse antibodies. Eur J Immunol 11:1012–6.

73. Nimmerjahn F, Ravetch JV. 2005. Divergent immunoglobulin g subclass activity through selective Fc receptor binding. Science 310:1510–1512.

74. Bruhns P. 2012. Properties of mouse and human IgG receptors and their contribution to disease models. Blood 119:5640–9.

75. Aschermann S, Lux A, Baerenwaldt A, Biburger M, Nimmerjahn F. 2010. The other side of immunoglobulin G: suppressor of inflammation. Clin Exp Immunol 160:161–7.

76. Chen X, Gula H, Pius T, Ou C, Gomozkova M, Wang LX, Schneewind O, Missiakas D. 2023. Immunoglobulin G subclasses confer protection against Staphylococcus aureus bloodstream dissemination through distinct mechanisms in mouse models. Proc Natl Acad Sci U S A 120:e2220765120.

77. Domanski PJ, Patel PR, Bayer AS, Zhang L, Hall AE, Syribeys PJ, Gorovits EL, Bryant D, Vernachio JH, Hutchins JT, Patti JM. 2005. Characterization of a humanized monoclonal antibody recognizing clumping factor A expressed by *Staphylococcus aureus*. Infect Immun 73:5229–5232.

78. Ganesh VK, Liang X, Geoghegan JA, Cohen ALV, Venugopalan N, Foster TJ, Hook M. 2016. Lessons from the Crystal Structure of the S. aureus Surface Protein Clumping Factor A in Complex With Tefibazumab, an Inhibiting Monoclonal Antibody. EBioMedicine 13:328–338.

79. Murphy AJ, Macdonald LE, Stevens S, Karow M, Dore AT, Pobursky K, Huang TT, Poueymirou WT, Esau L, Meola M, Mikulka W, Krueger P, Fairhurst J, Valenzuela DM, Papadopoulos N, Yancopoulos GD. 2014. Mice with megabase humanization of their immunoglobulin genes generate antibodies as efficiently as normal mice. Proc Natl Acad Sci U S A 111:5153–8.

80. Jefferis R. 2009. Glycosylation as a strategy to improve antibody-based therapeutics. Nat Rev Drug Discov 8:226–34.

81. Kim HK, Emolo C, Dedent AC, Falugi F, Missiakas DM, Schneewind O. 2012. Protein A-specific monoclonal antibodies and the prevention of Staphylococcus aureus disease in mice. Infect Immun doi:10.1128/IAI.00230-12.

82. Yu W, Kim HK, Rauch S, Schneewind O, Missiakas D. 2017. Pathogenic conversion of coagulase-negative staphylococci. Microbes Infect 19:101–109.

83. Claes J, Liesenborghs L, Lox M, Verhamme P, Vanassche T, Peetermans M. 2015. In Vitro and In Vivo Model to Study Bacterial Adhesion to the Vessel Wall Under Flow Conditions. J Vis Exp doi:10.3791/52862:e52862.

84. Bernardino PN, Bhattacharya M, Chen X, Jenkins J, Missiakas D, Thammavongsa V. 2023. A humanized monoclonal antibody targeting protein a promotes hopsonophagocytosis of Staphylococcus aureus in human umbilical cord blood. Vaccine 41:5079–5084.

